# Evolution and spread of SARS-CoV-2 likely to be affected by climate

**DOI:** 10.1101/2020.06.18.147074

**Authors:** Priyanka Bajaj, Prakash Chandra Arya

**Author notes:** Both authors have equally contributed to this work.

## Abstract

COVID-19 pandemic has been extensively studied by many researchers. However, it is still unclear why it was restricted to higher latitudes during the initial days and later cascaded in the tropics. Here, we analyzed 176 SARS-CoV-2 genomes across different latitudes and climate (Koppen’s climate) that provided insights about within species virus evolution and its relation to abiotic factors. Two genetically variant groups, named as G1 and G2 were identified, well defined by four mutations. The G1 group (ancestor), is mainly restricted to warm and moist, temperate climate (Koppen’s C climate) while its descendent G2 group surpasses the climatic restrictions of G1, initially cascading into neighboring cold climate (D) of higher latitudes and later into hot climate of the tropics (A). It appears that the gradation of temperate climate (Cfa-Cfb) to cold climate (Dfa-Dfb) drives the evolution of G1 into G2 variant group which later adapted to tropical climate (A) as well. It seems this virus followed inverse latitudinal gradient in the beginning due to its preference towards temperate (C) and cold climate (D). Our work elucidates virus evolutionary studies combined with climatic studies can provide crucial information about the pathogenesis and natural spreading pathways in such outbreaks which is hard to achieve through individual studies. Mutational insights gained may help design an efficacious vaccine.

**Graphical Abstract:** 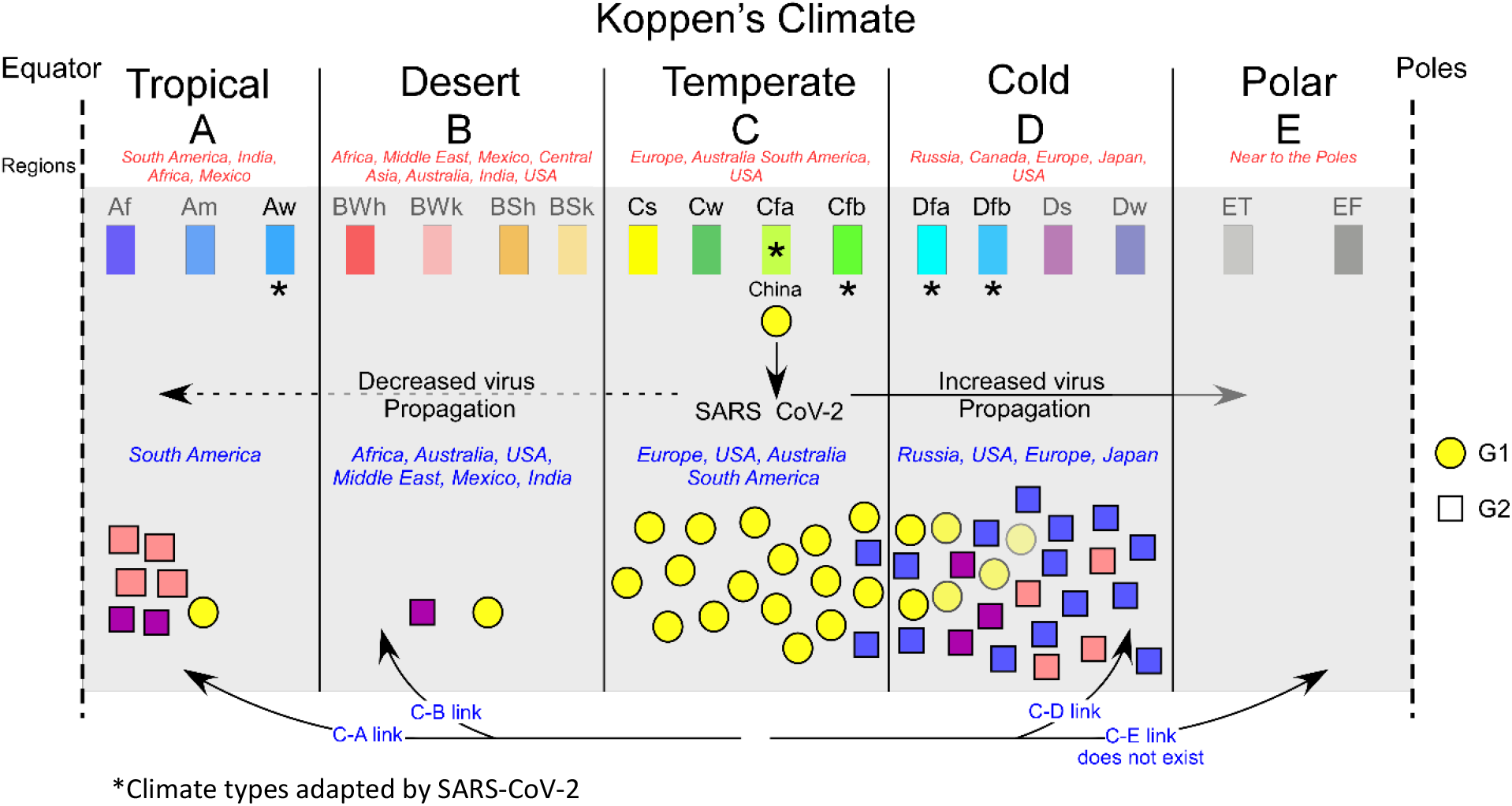

**In Brief:** The authors elucidate adaptation of SARS-CoV-2 to different climates by studying phylogenetics and the distribution of strains on Koppen’s climate map.

**Highlights:**

- Phylogenetic network divides SARS-CoV-2 strains into two variant groups, G1 and G2.
- G1 strains is restricted to Koppen’s *“temperate”* climate (mainly *Cfa-Cfb*).
- G2 strains has evolved from G1 to sustain in other climates mainly “*humid-continental” (Dfa-Dfb)* and “*tropical-savannah” (Aw)* climate.

## Introduction

The first case of Corona Virus Disease-19 (COVID-19) pandemic caused by (Severe Acute Respiratory Syndrome-2) SARS-CoV-2 pathogen was first reported from Wuhan, China^1^. In spite of various precautions such as lockdown, social distancing, wearing mask, and sanitization, the disease was able to reach almost every part of the world^2^. This zoonotic virus is like SARS (Severe Acute Respiratory Syndrome) coronavirus (79% similarity) and MERS (50% similarity) and is closely related to bat derived coronaviruses^1^. The SARS-CoV-2 can survive up to 3, 4, and 24 hours on aerosols, copper, and cardboard respectively, and up to 2-3 days on stainless-steel or plastic^1^ and spreads faster than its ancestors SARS-CoV-1 and Middle East Respiratory Syndrome (MERS-CoV)^1,3^. This led it concur a larger geographical region by infecting larger population. Since the social behaviour and travelling of humans have not changed much, what makes few respiratory viruses confined locally and others spread globally is still unclear. COVID-19 outbreak led to a big discussion, that does climate have a role in the spread of the disease? The ancestor SARS-CoV-1 losses its viability at higher temperature (38°C) and relatively higher humidity (>95%)^4^. Experiments support that SARS-CoV-2 is highly stable at 4°C but is sensitive to heat^5^. Studies both in favour and against have been published but the topic is still debatable^6,7^. A recent review compiles 61 studies relating COVID-19 with climate^1^. Several climatic factors such as humidity, precipitation, radiation, temperature, and wind speed affecting this virus spread have been incorporated. Both positive and negative association of COVID-19 with temperature and humidity have been published^1^. Recently, Carlson et al. mentioned that COVID-19 transmission could be affected by climate but discouraged the use of SDMs (Species Distribution Models) for COVID-19 transmission due to their limitations, which generally takes only climatic parameters as input, as they may not be appropriate tools^7^. This study was challenged by Araujo et al. mentioning strengths and limitations of the tools and reasoned that R_0_ of COVID-19 depends on several factors, it may also be affected by climate^6^. Since only climatic parameters are insufficient to capture climatic signatures of COVID-19 spread, such patterns can be recognized by combining phylogenetic and climatic studies^8^. Such approach enables to probe the similarities or differences in virus genome across similar or different climate types present all over the world. To understand such a behaviour, we have attempted to study the genomic sequence of the SARS-CoV-2 across different latitudes and climates.

Till now, pathogens are poorly mapped and little is known about their underlying ecological and evolutionary causes^9^. Nucleotide substitution has been proposed to be one of the most important mechanisms of viral evolution in nature^10^. However, factors responsible for the generation of these mutations are not well understood. One of the possible factors is adaptation to new environments, dictated by natural selection that discriminates among genetic variations and favours survival of the fittest^11^. Virus evolution as a consequence of climate change is poorly understood. SARS-CoV-2 consists of large single-stranded ~30kb long positive-sense RNA. These viruses majorly have a conserved genomic organization, consisting of a unique 265bp long leader sequence, ORF1ab polyprotein, and structural proteins like S (spike glycoprotein), E (Envelope), M (Membrane), and N (Nucleocapsid). ORF1ab encodes replicase, transcriptase and helicase, essential enzymes required for replication, along with non-structural and accessory proteins. Expression of non-structural proteins is facilitated by ribosomal frameshifting^12^. All coronaviruses express structural proteins S, E, M, N; spike glycoprotein being the most immunogenic to T-cell response^13^. Spike glycoprotein of coronaviruses binds to human angiotensin-converting enzyme 2 (hACE2) receptor for viral fusion and entry and is the main target for neutralizing antibodies and development of vaccines^14^. Membrane protein is also antigenic as it stimulates a humoral immune response^15^. E protein is responsible for virus assembly and release of virion particles^16^. Nucleocapsid protein packages RNA genome into a helical ribonucleocapsid protein (RNP) complex during virion assembly and is capable of eliciting an immune response^17^. Since it is still not clear whether SARS-CoV-2 evolution and spread have relation with climate, our study may act as a missing link between genomic sequence, climate and COVID-19 severity. If SARS-CoV-2 is responding towards external climate it can be delineated by superimposing its genomic variants across different latitudes and Koppen’s climate^8^. The earliest and the most simple classification of Earth’s climate is based on latitudes which divide the Earth’s climate into seven climate zones, North Frigid Zone (NFZ), North Temperate Zone (NTZ), North Subtropical Zone (NSTZ), Tropical Zone (TZ), South Subtropical Zone (SSTZ), South Temperate Zone (STZ) and South Frigid Zone (SFZ) lying between 90°N to 66.5°N, 66.5°N to 30°N, 30°N to 23.5°N, 23.5°N to 23.5°S, 23.5°S to 30°S, 30°S to 66.5°S and 66.5°S to 90°S, respectively^18^. Based on temperature and precipitation, Wladimir Koppen divided Earth’s climate into five major climates, A (Tropical), B (Arid), C (Temperate), D (Cold or Continental) and E (Polar) which are further subdivided into 30 climate types^19^. To understand the effect of climate on SARS-CoV-2 evolution, the present study comprises of two parts: (1) Sequence analysis of SARS-CoV-2 strains, (2) Mapping SARS-CoV-2 strains across different climates. These combined studies can provide insights on within species evolution and preferential distribution of SARS-CoV-2 across different climatic zones which might be difficult to probe through individual studies.

## Methodology

### Molecular phylogenetic analysis

Approximately, 11,000 full-length genome sequences of SARS-CoV-2 were available in Global Initiative on Sharing Avian Influenza Data (GISAID) database, accessed till 2^nd^ May 2020. 185 full-length SARS-CoV-2 genomic sequences from countries across the globe, with genome length more than 29 kb and high coverage were obtained from GISAID database and the reference genome was retrieved from GenBank24 (Supplementary Table S1). This sample size was selected by taking 95% confidence interval, 0.5 standard deviation and 7% margin of error. To avoid bias related to the geographical area covered by a country, genomic sequence of strains isolated from different locations from each country/ climate type was retrieved, depending on the availability of data. The corresponding location, latitude, Koppen’s climate, Koppen’s Climate type, SARS-CoV-2 variant group, Environment/ Region, Climate zone, temperature, and precipitation of each strain is provided in Supplementary Table S2. These sequences were aligned to the full reference genome^20^ by using Biomanager and Seqinr packages of R (version 3.6.3). Among 185 genomes, some partial genomes were discarded. Thus, finally 176 genomes were analysed. NC_045512 genome sequence was used as reference and the genomic coordinate in this study is based on this reference genome. Based on protein annotations, nucleotide level variants were converted into amino acid codon variants for alignments when its location within a gene was identified. The amino acid position numbering is according to its position within the specified gene (Coding Sequence) as annotated in reference sequence (NC_045512, NCBI)^20^. To ensure comparability, we trimmed the flanks of all sequences. The aligned sequences were used to construct a phylogenetic tree using MEGA X^21^. The evolutionary history was inferred using the Neighbor-Joining method (500 bootstrap tests)^22^. The optimal tree with the sum of branch length = 0.01116462 is shown. The tree is drawn to scale, with branch lengths in the same units as those of the evolutionary distances used to infer the phylogenetic tree. The evolutionary distances were computed using the Maximum Composite Likelihood method^23^ and are in the units of the number of base substitutions per site. All ambiguous positions were removed for each sequence pair (pairwise deletion option). A total of 29408 positions were present in the final dataset. The results are presented in the form of DNA sequencing i.e., U (uracil) is read as T (thymine). We have labeled each virus strain by the GISAID Accession ID and the location from which it was isolated in the format “Location|EPI ISL Accession ID”, in the constructed phylogenetic tree. For ease of visualization, we have marked a new Strain ID (1 to 176) against each SARS-CoV-2 isolate in the phylogenetic tree (Figure 1). The same Strain ID is used for the climatic studies in this article. High-frequency SNPs (Single Nucleotide Polymorphisms) distinguishing one virus cluster from the others is referred to as “virus cluster SNPs” throughout this paper.

**Figure 1:**
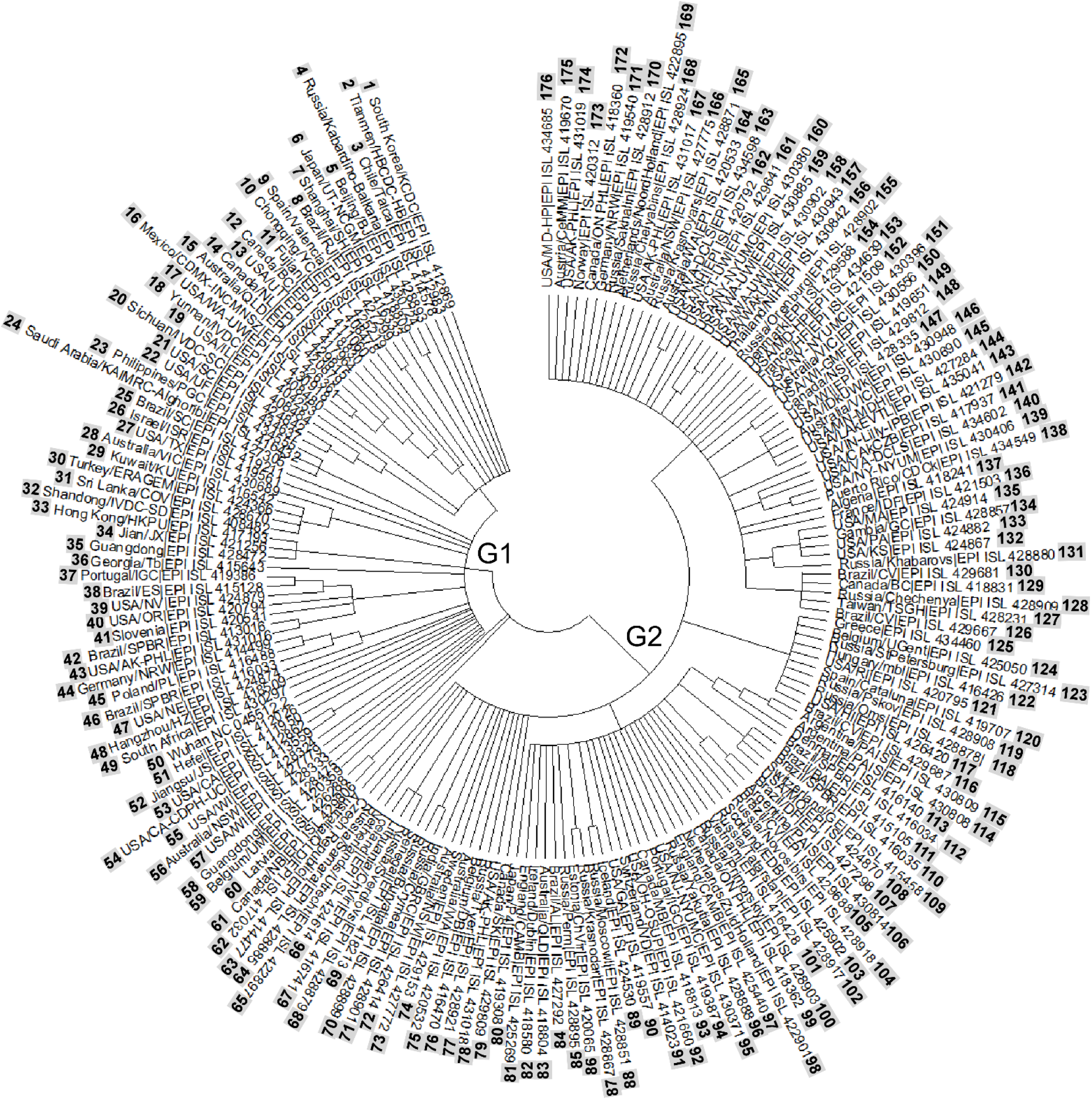
Phylogenetic network divides 176 SARS-CoV-2 strains into two variant groups. Largely, the left side of the tree (1 to 58) constitute the G1 group and the right side of the tree constitutes the G2 group (59 to 176). Branch length is proportional to the genomic relatedness of the viral isolates. Closely related virus isolates comprise the same SNP with respect to the reference genome (Strain ID: 50) and form a cluster. The evolutionary history of 176 taxa was inferred using the Neighbor-Joining method^22^ (500 bootstrap tests). A total of 29408 positions were analysed with nucleotide position numbering according to the reference sequence^20^.

### Mapping virus strain on the Koppen’s climate map

The location of each SARS-CoV-2 strain is obtained from the METADATA file provided in GISAID database for each viral isolate (Supplementary Table S1). The coordinates of the locations were taken from the official website of USGS Earth Explorer^24^. The Gieger-Koppen’s climate map is used for climatic studies^19^. The Koppen’s climate type, temperature, precipitation of each strain is assessed from weatherbase^25^ and CLIMATE.ORG^26^. The above information for each strain is tabulated in Supplementary Table S2. The map is georeferenced by using ‘Arc-GIS 10.1’^27^. The locations of all strains (n=176) were transferred to the georeferenced map^27^. On the map, the G1 strains were symbolized as ‘Yellow-circle’, and G2 as ‘Square’ (Figure 3). Each strain in the map is labelled as per their Strain ID (1 to 176) (Figure 3), the map combines information of the phylogeny, climate, and global distribution of SARS-CoV-2. These locations were classified into coastal and continental region, we define the coastal region as land region < 500 km from the ocean/sea and the continental region as land lying >500 km from the coastline measured through google maps.

### Statistical analysis

Chi-square were performed in Microsoft Excel (2016) to test our null hypothesis. Null hypothesis for these tests are mentioned in the text and its corresponding contingency table is mentioned in Supplementary Table S2. Histograms depicting the distribution of coronavirus in coastal region, continental region, Koppen’s climate and climate type were plotted using R (version 3.6.3). SigmaPlot10 was used to generate box plot, regression plot, and mesh plot to statistically compare frequency distribution of latitude, temperature, and precipitation of G1 and G2 strains. We performed one-way ANOVA to estimate statistical differences in the latitude, temperature and precipitation between G1 and G2 virus populations. Various scatterplots between latitude, temperature, and precipitation of G1 and G2 strains were plotted in R (version 3.6.3). Values were considered statistically significant for P values below 0.05. Exact P values are provided in appropriate Figures.

### Data accessibility

The full-length genomic sequences were downloaded from GISAID website (https://www.gisaid.org/), an open-source database for influenza viruses. The data is downloaded as FASTA file along with the acknowledgement. The location of each strain is accessed from its METADATA file. The Koppen’s Climate map is taken from the published article^19^. The Koppen’s climate type, temperature and precipitation for each strain is taken from weatherbase (https://www.weatherbase.com/) and CLIMATEDATA.ORG (https://en.climate-data.org/). Refer Supplementary Tables S1-S5. The code is available from the corresponding authors on request.

## Results

### Molecular phylogeny analysis to infer genomic similarities and their distribution in different climates

To probe genomic similarities between SARS-CoV-2 virus isolates, a phylogenetic tree was constructed by aligning 176 virus genomes to the reference genome^20^ retrieved from GISAID. Interestingly, our Multiple Sequence Alignment (MSA) results reveal sixty virus cluster Single Nucleotide Polymorphisms (SNPs) (see methodology). Table1 comprises of SNPs of virus clusters across different climatic zones, Koppen’s climate and climate type. Climatic parameters (temperature and precipitation) for each virus strain is mentioned in Supplementary Table S2. Based on phylogenetic clustering, 176 SARS-CoV-2 strains are majorly divided into two groups, we named them as G1 (1-58) and G2 (59-176) (Figure 1). Predominantly four mutations distinguish G2 from G1 group, i.e., 1) a synonymous mutation (C241T) appeared in the unique leader sequence, 2) F924 (C3037T) appeared in nsp3, encoding for papain-like proteinase^28^, 3) a non-synonymous mutation, P214L (C14408T) arose in ORF1b, that codes for four putative non-structural proteins (nsp13, nsp14, nsp15 and nsp16), functionally involved in replication-transcription complex^29^, and 4) D614G (A23403G) arose in S gene, encoding spike glycoprotein^13^ (Figure 2a). Among four mutations in G2, the D614G mutation, lying in spike glycoprotein was widely studied due to its higher infectivity and involvement in entering the host cell through hACE2 receptors^30–33^. The other three mutations in G2 have co-evolved with D614G making it distinguishable from G1. We explored the extent of genome-wide divergence of G1 and G2 group across different climate zones and Koppen’s climate (Figure 2b). 59% of G1 viruses fall in NTZ, 14% in NSTZ, 12% in TZ, 10% in SSTZ and 5% in STZ. 76% of the virus isolates in G2 group are present in the NTZ, 13.5% in TZ, 7.6% in STZ and remaining 2% is equally distributed in NSTZ and SSTZ, showing G2 strain variants evolved to adapt to temperate zones as their population decreased drastically in the subtropical zones. These results show both G1 and G2 strains have a strong preference towards higher latitudes i.e., NTZ (Figure 2c). Mapping viral strains on Koppen’s map (thoroughly discussed in the next section) reveal their prevalence majorly in the C and D climate (Figure 2d). 71% of G1 lie in C climate, 17% in D and the remaining is equally distributed in the A and B climate. 54% of G2 lie in C climate, 36% in D, 9% in A and 1% in B climate pointing towards a preferential shift of the novel coronavirus towards D climate (Figure 2b), alluding G2 is climatically and genomically more diverse than G1. The analysis suggests that the G1 group is mostly restricted to temperate climate (C) and G2 is climatically and geographically widely distributed, it is possible that these mutations were acquired by G1 to stabilize itself in different climates hence allowing it to spread globally. Similar climatic concordance with the temperate climate (C) was also observed for SARS-CoV-1 that was responsible for 2002-2004 epidemic as it prevailed in regions of Australia, Europe, Canada and, China^34^, having Koppen’s C climate. Such similar occurrence of SARS-CoV-1 and G1 group of SARS-CoV-2 hints towards, why initially G1 variant group (consisting of the reference genome NC_045512^20^) that has 79% similarity to SARS-CoV-1^1^ was majorly located in the temperate climate (C), and latter it evolved into G2 variant group that allowed it to extend its climatic boundaries into temperate, cold and tropical climate. These results suggest these four SNPs could be the key factors in increasing the virulence, transmission and sustainability of the virus in humans.

**Figure 2:**
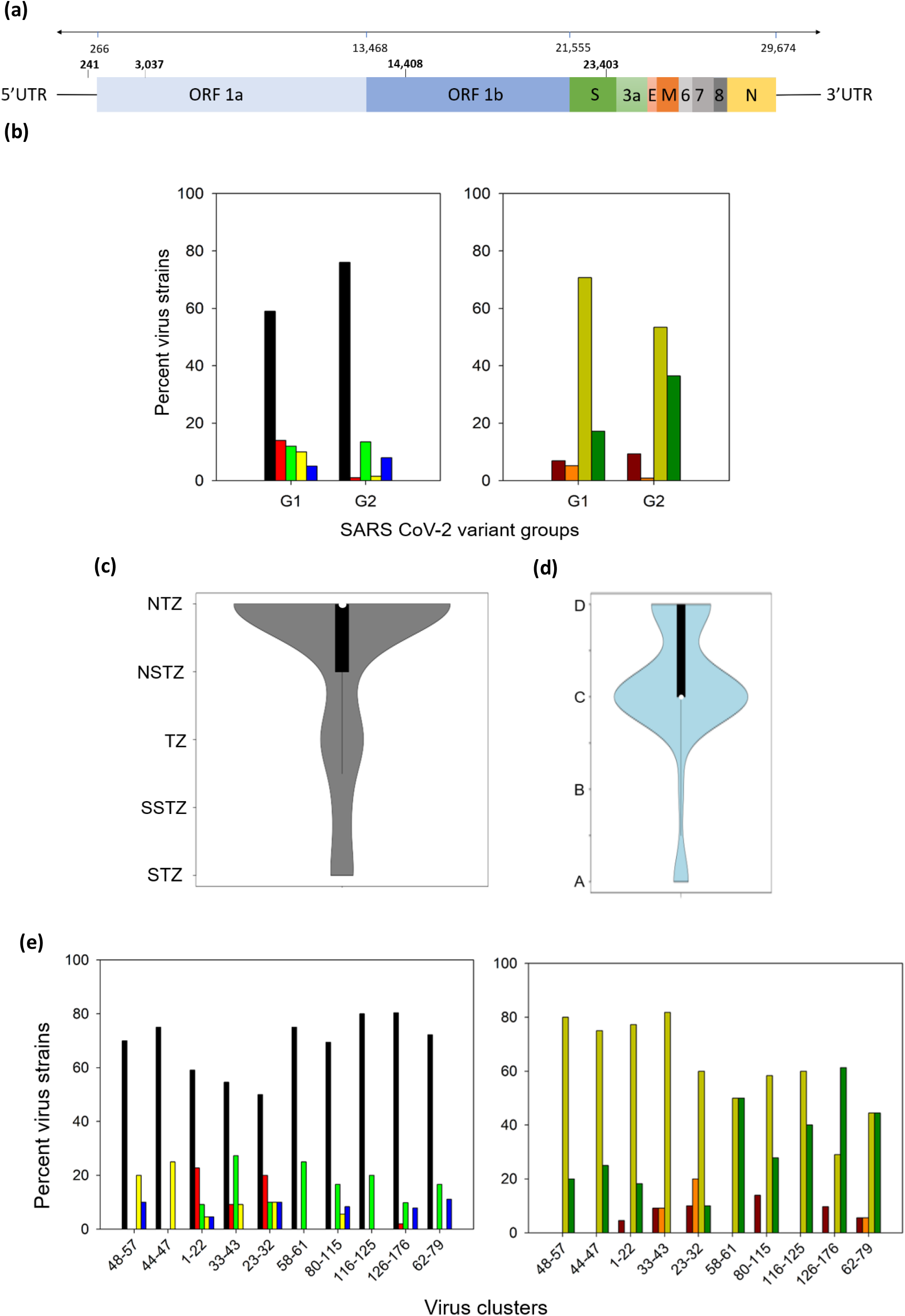
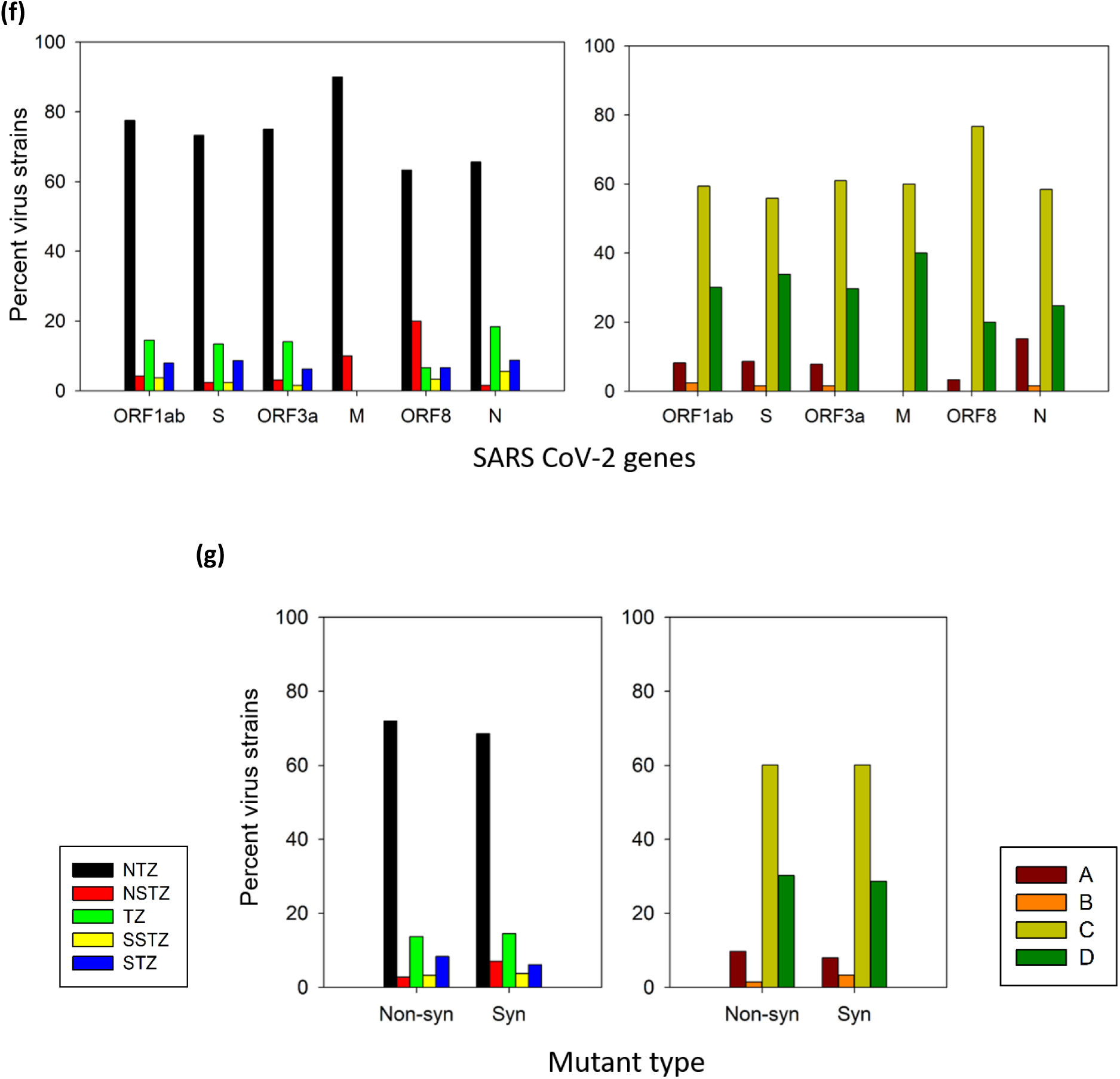
Molecular phylogeny analysis to infer genomic similarities of SARS-CoV-2 and their distribution across different climate zones^18^ and Koppen’s climate types^19^. (a) Genomic architecture of SARS-CoV-2 genome highlighting four positions, substitutions on these positions probably enabled evolution of G1 into G2 variant group. (b, e-g) Strains found within a virus cluster (as shown in the phylogenetic tree and mentioned in Table 1) were analysed for significant mutations that may have arisen due to climatic pressure. Hence, percentage of such virus strains is plotted according to the geographical location of the climate zone from where they were isolated. The height of the bar is proportional to percent virus strain occurring in the specified condition i.e., labelled on the x-axis. Box in the left panel consist of color code for each climate zone and box in the right panel consist of color code for Koppen’s climate. Left panel shows distribution of percent virus strains in different climate zones and right panel shows distribution of percent virus strain in Koppen’s climate (b) Percent virus strains prevailing in different climate zones, stratified by SARS-CoV-2 variant groups. Width of curves of violin plot is proportional to the number of SARS-CoV-2 strains (n=176) in varied (c) climate zones and (d) Koppen’s climate. (e) Abiotic factors influencing evolutionary dynamics of phylogenetic virus clusters. (f) Percent of virus strains with high frequency SNPs in each gene. (g) Type of mutation i.e., non-synonymous (Non-syn) or synonymous (Syn) exhibited by viruses.

We further analyzed the order in which the phylogenetic clusters evolved from the ancestor 45-57 cluster (containing the reference genome, Strain ID: 50) based on nodes, mutational branches and branch length. The order in which the virus evolved is 44-47 (G1440A, G392D; G2891A, A876T), 1-22 (C8782T, S2839; T28144C, L84S), 33-43 (G26144T, G251V), 23-32, 58-61 (C15324T, N519), 80-115 (G28881A, G28882A, R203K; G28883C, G204R), 116-125 (A20268G, L2167), 126-176 (G25563T, Q57H) and 62-79 (cluster, acquired genomic mutation and its corresponding amino acid mutation). In Figure 2e, looking at the distribution of the viruses in different climate zones, no such preference was observed as the virus evolved. Virus cluster 58-61, linking G1 and G2 has an equal distribution of virus strains in C and D climate. The virus cluster 80-115 of G2 more closely related to G1, is widely distributed in A, C and D climates. Within 80-115 virus cluster, 106-115 cluster shows distribution in C and A climate. A trend was observed that virus clusters in G2 group gradually evolved to sustain in Koppen’s D climate (80-115 to 116-125 to 126-176). Within these major virus clusters, small clusters also exist as shown in Table1 with their mutations along with their climatic distribution.

We have examined whether climatic conditions exhibit any selective pressure on each gene (Figure 2f). Since, the present picture of the data shows that SARS-CoV-2 spreads widely in NTZ, as expected all genes are having mutations in NTZ, suggesting the virus is probably using varied mechanisms to adapt to the two main climates of NTZ i.e., temperate (C) and cold or continental (D). Mutations in the M gene are only pertaining to NTZ and NSTZ and are present in C and D climate. In particular, there is a surge in the virus strains carrying SNPs in ORF8 in the NSTZ (20%). 77% of the SNPs in ORF8 lie in the C and 20% in the D climate. Overall, the distribution of virus cluster SNPs of ORF1ab, S, ORF3a, and N gene follows a similar pattern across all the climatic zones and Koppen’s climate, implying no difference in selective pressure of the climate in generating mutations in these genes. S, M, and N proteins are immunogenic^13,15,17^, implicating virus evades immune response by introducing these substitutions.

Apart from non-synonymous mutations, synonymous mutations within the gene can also significantly affect protein function due to codon usage bias^35^ and through mechanisms such as ribosome stalling^36^ and mRNA secondary structure formation^37^. We probed the frequency of derived synonymous versus non-synonymous mutations and observed a very similar distribution pattern of the derived synonymous versus missense mutations across all climate zones and Koppen’s climate (Figure 2g). These analyses suggest novel coronavirus is using varied mechanisms both at the transcriptional as well as translational level to adapt, survive, and increase infectivity in all types of climates. These findings unequivocally bolster a requirement for further prompt, comprehensive studies that join genomic information, epidemiological information, and climatic distribution with COVID-19 severity.

### Distribution of strains across Koppen’s climate

To probe the relation between climate and SARS-CoV-2 strains, we superimposed genomic information along with their geolocations on the climate map of Wladimir Koppen (Figure 3). We carefully examined the distribution of strains on Koppen’s map and an overview of the map shows, the distribution of 176 strains are mainly concentrated in the western coasts of Europe and North America, and eastern coasts of China, North America, Australia and South America (Figure 3). Throughout the text Koppen’s climate type is marked within quotations and its standard symbol is written within brackets e.g., “*humid-subtropical” (Cfa)*. List of Koppen’s symbol of each climate type is given in Supplementary Table S3 and its criteria for classification is given in Supplementary Table S4. Mostly the SARS-CoV-2 strains are distributed in the *“humid-subtropical” (Cfa)* and *“marine-temperate” (Cfb)* and *“humid-continental” (Dfa-Dfb)* climate and, two strains from virus clusters (80-115 and 126-176) belonging to South America, are found in “tropical-savanna” *(Aw)* of ‘A’ climate (Supplementary Table S5). The map displays ~86% (151/176) of virus isolates are distributed in the coastal regions and the remaining in continental region (Chi-square test, *P*<.001 considering null hypothesis that equal number of virus isolates are found in coastal and continental region, Figure 4a, Supplementary Table S6). Around ~74% (130/176) of the total strains are distributed in “humid-subtropical” (Cfa) and “marine-temperate” (Cfb) climate type of C climate and “humid-continental” (Dfa-Dfb) climate type of D climate. The remaining ~26% (46/176) strains are distributed in other climate types of Koppen’s climate including non ‘Cfa-Cfb’ of C climate and non ‘Dfa-Dfb’ of D climate (Figure 4b). It seems that spread of COVID-19 is maximally in areas with ‘Cfa’ and ‘Cfb’ climate type. The climatic parameters (temperature and precipitation) in which these strains lie, were analyzed. Statistically, significant difference was found in the latitudes of G1 ~24.14±3.5 (mean±s.e.) and G2 ~34.03±2.7 (one-way ANOVA, *P*=.03251, Figure 5a). Statistically, significant difference was observed in the temperatures of G1 (15.82±0.75 °C (mean±s.e.) and G2 (11.67±0.68 °C) strains (one-way ANOVA, *P*<.001, Figure 5b). However, the difference in precipitation for G1 (1046.95±80 mm) and G2 (896.64±35.48 mm) strains is statistically not significant (one-way ANOVA, *P*=.06118, Figure 5c). The latitudes and temperature are inversely related to each other (r = −0.6649, Supplementary Figure S1a), which explains the occurrence of G1 strains in lower and G2 strains in higher latitudes (Figure 5d). Such relation between latitude and precipitation has not been observed (r = −0.3064, Supplementary Figure S1b and Figure 5e). A mesh plot simultaneously evaluates all climatic parameters for both G1 and G2 strains, the results agree to the limited temperature and wider precipitation range of G1 group and interestingly the G2 group appears in a wider temperature and shows a preferential shift towards lower temperature which is evident from the fact that it initially appeared more in higher latitudes (Supplementary Figure S2). A complete description of the distribution of G1 and G2 strains lying in different countries and/or continents of the world is provided in Supplementary Figure S3 and Supplementary Material.

**Figure 3:**
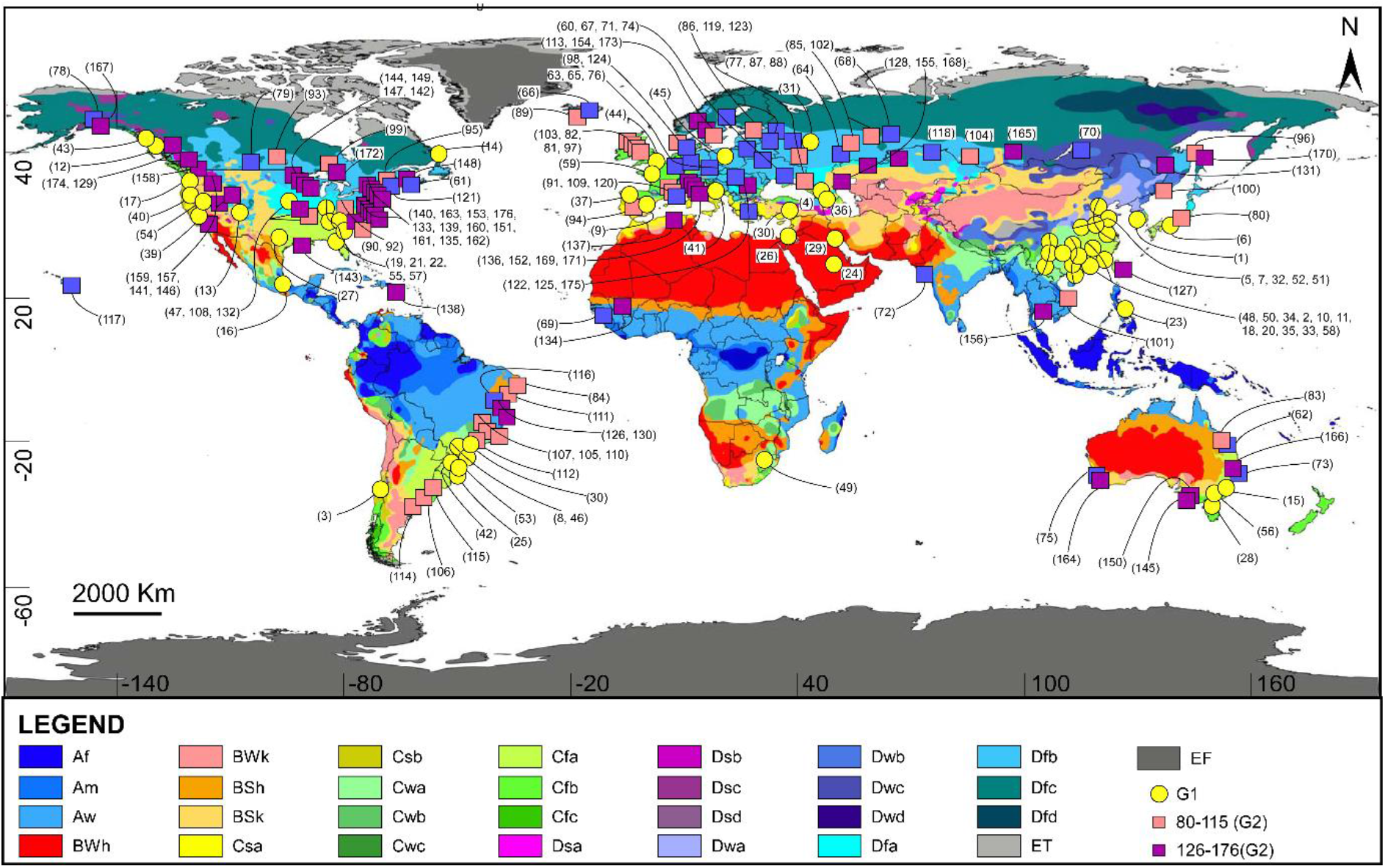
Global distribution of SARS-CoV-2 strains on the Gieger-Koppen’s map displaying different climate types^19^. Each strain is labelled as per the strain ID (1 to 176) within parenthesis. The G1 strains were symbolized as ‘Yellow-circle’, and G2 as ‘Square’, pink square denotes strain clusters (80-115) stable across C, D and A climate, purple square represents strain cluster (126-176) stable majorly in D climate, the remaining G2 strains (blue squares) are stable across C and D climate. Standard Koppen’s climate-type symbols are mentioned in the legend, the criteria for distinguishing these climate types, is mentioned in Table S3. Table S4 contains full form of these symbols. All symbols with initials ‘A’ (Af, Am, Aw) are of tropical climate, initials with ‘B’ belong to desert climate, ‘C’ to temperate and ‘D’ to cold and ‘E’ to polar climate. The shades of blue on the map, in North America and Russia belongs to D climate. The shades of blue in South America, Africa, and South Asia belong to tropical climate. Shades of yellow and green belongs to C climate. Shades of red, orange and pink belong to Desert climate.

**Figure 4:**
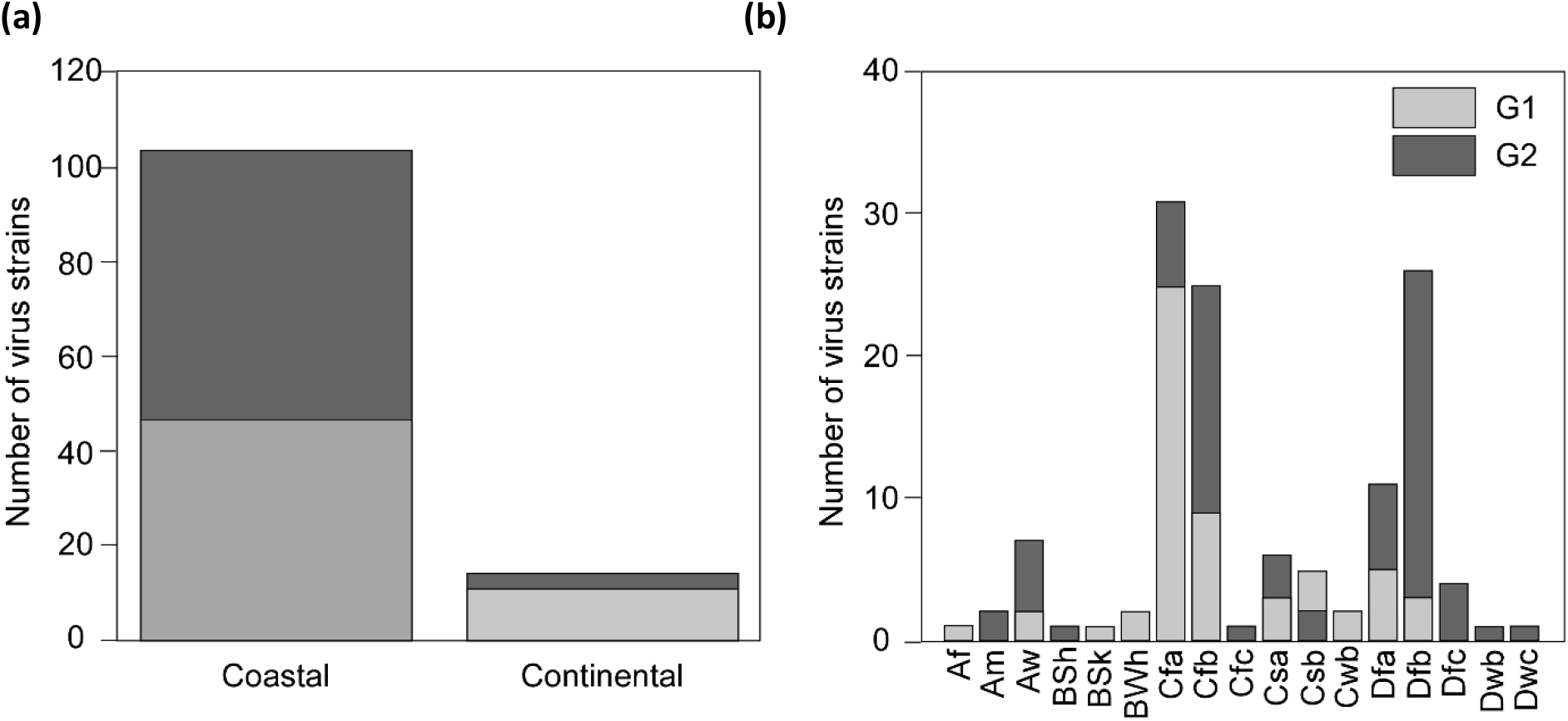
Global distribution of SARS-CoV-2 strains (n=176) (a) in the coastal and continental region (b) and in different Koppen’s climate types^19^. Number of virus strains in G1 population is represented by light grey color and of virus strains in G2 population is represented by dark grey color.

**Figure 5:**
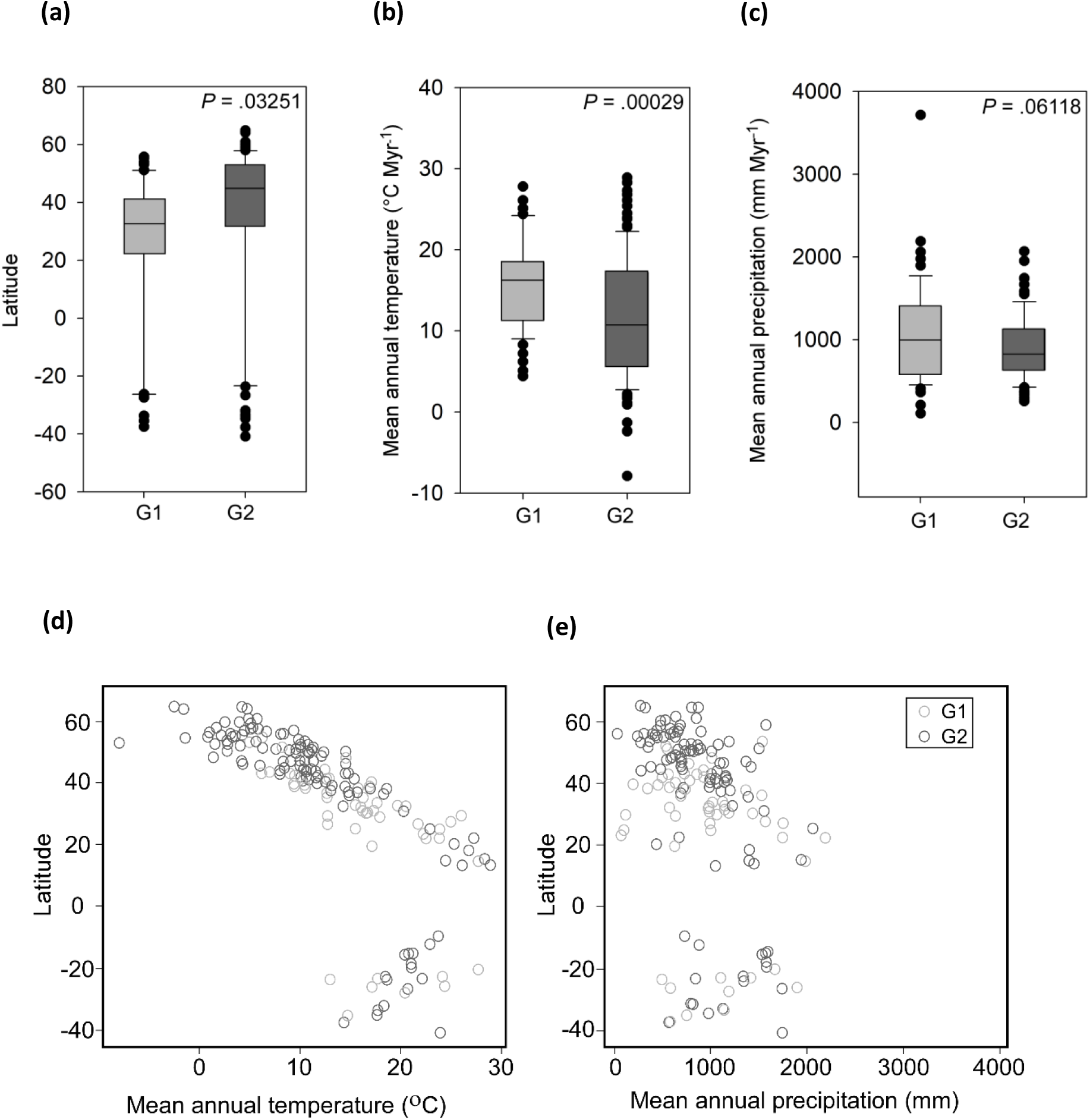
Comparative analysis of different climatic parameters such as latitude, temperature and precipitation between G1 and G2 variant groups. (a) Positive values represent the latitude range falling in Northern Hemisphere and negative values represent latitude range falling in Southern Hemisphere. The G2 strains preferentially occur towards the higher latitudes than G1 (*P*=.032; 95% CI 17.12-31.12 for G1; 95% CI 28.67-68.06 for G2). (b) The mean annual temperature of G2 is significantly lower than the G1 strains (*P*<.001; 95% CI 17.32-14.32 for G1; 95% CI 13.02-10.33 for G2) (c) Mean annual precipitation of G1 and G2 strains is nearly same (*P*=.061; 95% CI 1207.16-886.75 for G1; 95% CI 966.91-826.37 for G2). (a-c) Black horizontal line in the middle of the box is median, upper and lower limits of the box indicate first and third quartile. Black dots represent outliers. P values is based on one-way ANOVA. Scatter plot for (d) latitude and annual temperature and (e) latitude and precipitation for each SARS-CoV-2 strain (n=176) belonging to G1 group (n=58, shown in light grey) and G2 group (n=118, shown in dark grey).

## Discussion

A pattern observed and discussed by several authors is that SARS-CoV-2 predominantly affected higher latitude countries (eg., China, Europe, USA and Australia, Japan, Turkey etc.) in its preliminary stage (November 2019 to March 2020). This led many researchers predict a reduced spread in regions with higher temperature such as tropical countries^38,39^. Our phylogenetic analysis provides one more dimension to the above observations in which we have identified two groups G1 and G2, similar observations were also reflected in other studies^40,41^. We have further integrated the genomic information with climatic data and analyzed the spread of these two variant groups across the globe and its climate.

There can be many reasons behind the observed pattern of spread. First, it could be because of less testing in tropical countries. However, if this would be true, local media would have reported over-crowding of patients in hospitals. For example, Italy escalated its testing when media reported over-crowding of hospitals with patients having COVID symptoms. But such incidences were not reported from tropical countries. Second, it could be higher human mobility between China US, and China Europe. But mobility between China and South-East Asia is also very high. Thus, no outbreak in tropical countries initially is perplexing (till March 2020). Third, most countries implemented lock-down measures from mid-March. Until March people were working in normal mode. Hence, it is not possible that stringent measures in containing the virus evaded it from cascading into the tropical countries. Fourth, the population density of the tropics could be lower than temperate countries. But this is also not true, in fact most of world population resides in tropical countries and are mostly developing with limited proper health and hygiene facilities compared to temperate countries. In such situation the disease should have equally spread in tropics like that of temperate countries as positive cases of COVID-19 were identified from many tropical countries in January itself. Fifth, our analyses show that G1 group spread largely in temperate climate. A possible reason behind this could be that host-immune response of people residing in the tropical countries resisted the infection. However, no supporting evidence have been found till now and needs detailed investigation.

Hence, these reasons negate the fact that the observed patterns could solely be due to community structure, social dynamics, government policies, global connectivity, population density and number of reported cases. Similar observations i.e. more COVID cases in higher latitudes were reflected in COVID-19 growth curves of Yusuf and Bukhari until March 2020 for different countries, which led them interpret that natural factors (temperature and humidity) are responsible for less number of COVID cases in tropical countries^42^. Thus, we are inclined to say that environmental factors, directly or indirectly play a role to explain the observed pattern of spread in cold countries during the initial stages. Since, in winters mostly people stay indoors, thus climate can indirectly be involved in increased growth rate of virus in cold countries. However, if climate indirectly had a role to play in the spread, spread of the disease would have been limited in summers in tropical countries. On the contrary, the COVID-19 outbreak in tropical countries started in summers, soon after emergence of G2 strains.

Results of this study depict that in the initial stages (until March 2020) of the pandemic the G1 group was restricted to temperate climate (Koppen’s C type) whereas G2 group spread all over the world. Our analyses suggest that same regions were affected by G1 variant group and SARS-CoV-1. It is known that SARS-CoV-1 was not able to survive in regions with warm climate as its lipid bilayer was prone to degradation at higher temperature^43^ and it loses its viability at higher temperature^4^. A recent study shows the structural stability of SARS-CoV-2 Virus Like Particles (VLPs) degrades on increasing temperature^44^, supporting our observations.

Our analyses favor that evolution of G1 into G2 helps to sustain this virus from temperate to cold and tropical climate successively, mainly due to four mutations i.e., in leader sequence, ORF1ab, and S gene. The leader sequence and ORF1ab is involved in replication and transcription, and the S gene is involved in binding to the host cell through hACE2 receptors. Substitutions in the ORF1ab gene may increase the synthesis of replicase-transcriptase complex, thus, increasing the replication rate of the virus and blocking the host innate-immune response. 614 position in spike glycoprotein lies near the S1/S2 subunit junction where the furin-cleavage site is present (R667) that enhances virion cell-cell fusion^45^. This suggests, aspartate to glycine substitution in the vicinity of the furin-recognition site may result in a conformational change of the spike glycoprotein that favors higher affinity of the Receptor Binding Domain (RBD) to hACE2. A recent article showed retroviruses pseudotyped with Glycine at 614 position infected ACE2-expressing cells markedly more efficiently than those with Aspartic acid due to less S1 shedding and greater incorporation of the S protein into the pseudovirion^30^. Several studies reported D614G mutation is increasing at an alarming rate^31,32^. Few observed that this alteration correlated with increased viral loads in COVID-19 patients^31^. This is consistent with the epidemiological data showing proportion of viruses bearing G614 is correlated to increased case fatality rate on a country by country basis^33^. This substitution coevolved with substitution in the leader sequence, nsp3 and RdRp (RNA dependent RNA polymerase) proteins, suggesting these mutations allow the virus to transmit more efficiently. This explains these mutations have not emerged merely because of founder’s effect but this virus under selection pressure has made itself more stable and infective. Also, Forster et al. observed in his phylogenetic analysis, the preferential geographical spread of SARS-CoV-2 and provided a plausible cause which could be founders effect or immunological or environmental effect^41^. Although there is a possibility that the stable variant might have appeared because of host innate immune response or some unknown reason, in such a case it would not show any close association with climate. Through our analysis, we are inclined to say that climate has an effect of SARS-CoV-2 evolution. However, in particular, selective pressure of climate on each gene of SARS-CoV-2 is not visible. Our genomic analysis of virus strains show that the novel coronavirus undergoes both synonymous as well as non-synonymous mutations throughout its genome in various climates, suggesting the novel coronavirus uses multiple mechanisms both at the transcriptional and translational level for evading the immune response, developing drug resistance and increasing pathogenesis. However, the actual role of these mutations is not yet determined, and these studies need to be further enlightened by biophysical and biochemical studies. Such mutational insights will aid the design of efficacious vaccines that can be stored and transported in a wide range of temperature and conditions, thereby minimizing cold storage costs.

To delineate the signatures of underlying abiotic factors (temperature, precipitation, and latitude) responsible for evolution of SARS-CoV-2 (n=176), spreading patterns of G1 and G2 strains were carefully examined on Koppen’s climate map. Figure 3 shows an elevated spread of COVID-19 in the western and eastern coasts of the continents and a diminished spread in the hot and cold deserts. The G1 strains are majorly present in the eastern and western coasts of the continents and G2 strains lie in both the coastal regions and continent’s interior. On a closer inspection, the eastern coasts of continents consist of “humid-subtropical” (Cfa) climate while the western coasts of continents consist of “marine-temperate” (Cfb), commonly known as east and west coast climate, respectively. These two climates are very similar to each other and belongs to temperate climate also known as C type climate of Koppen’s classification scheme. A very large portion (~94%) of habitable China consists of temperate climate (C), i.e., humid subtropical climate (Cfa), which explains presence of only G1 strains in China and one strain of G1 is present in cold climate (D) present near the transition of temperate (C) to cold climate (D), thus probably temperate climate was suitable for G1. A similar association of G1 with temperate climate (C) was found in eastern and western coast of North America, eastern coast of South America, western coast of Europe and eastern and western coast of Australia. Statistically, distribution of G1 strains all over the globe is in concordance with the temperate climate and strongly favor C climate (Chi-square test, *P*<.01 for null hypothesis that G1 strains are equally distributed in all climates, Supplementary Table S6) as compared to any other climate. If climate does not have any role in the evolution and preferential spread of coronavirus, in such a case G1 would have been evenly distributed in all climate types which is not the case. Few exceptions of G1 seen in other climate types are most probably because of travel as they remained subsided in that climate, implying their inability to sustain in other climate types. It appears that the G1 strains existed in temperate climate all over the world but could not extend their geographical territories beyond temperate climate. Contrastingly, the evolved G2 strains can sustain in temperate (C), cold (D) and tropical (A) climate surpassing the climatic restrictions of G1. Map interpretation suggests that G2 strains enters the continent’s interior through D climate (e.g., North America and Russia). Temperate climate (C) generally grades into cold climate (D) and deserts (B) in the northern hemisphere (e.g., C to D: Europe to Russia, and USA to Canada; C to B: China, and USA). In southern hemisphere, gradation of temperate climate (C) into tropical climate (A) and deserts (B) exists (e.g., C to A, Brazil; C to B, Australia), C to A transition is identified by virus cluster 105-115 in phylogenetic tree. In Russia, 91.3% (21/23) of the strains belong to G2 (Figure 3), are mainly present in the ~8500 km long and 600-1700 km wide D climate belt (‘Dfa-Dfb-Dw’), suggesting the G2 strains might have adapted to the D climate (Chi-square test, *P*<.001 for null hypothesis that both G1 and G2 strains are equally found in D climate of Russia, Supplementary Table S6). Similar observations are seen for North America, South America, and Australia. The eastern and western coasts of North America have temperate climate and are connected by cold climate along USA-Canada boundary (i.e., having humid subtropical (Cfa) in eastern coast and marine temperate climate (Cfb) in western coast) (Figure 3). The G2 strains follows this cold climate (Dfa-Dfb) belt which is ~3800 km long and ~600 to 1000 km wide. The dominance of G2 and nearly absence of G1 population in cold climate of North America is similar to the observations of Russia. Our analyses suggests that a fall of temperature from temperate to cold climate might have dictated the evolution of G1 into G2 variant group. Similarly, a change in climate from C to A probably made the strains stable in tropical regions. Overall, our analyses suggest that SARS-CoV-2 has likely evolved to sustain in different climate, thereby increasing its spread. Studies combining genetic information with climate can provides useful information about virus evolution and possible climatic pathways during an outbreak.

## Conclusion

It is reasonable to assume COVID-19 transmission pathway and evolution is influenced by climate. Phylogenetic network classified 176 SARS-CoV-2 strains into two variant groups G1 and G2. The G1 strains were habituated to C climate that evolved into G2 by undergoing significant mutations (C241T in leader sequence, F924 in ORF1a, P214L in ORF1b and D614G in S gene), plausibly extended its climatic boundaries from C to D climate, displaying role of natural selection on virus evolution. In our analysis SARS-CoV-2, were found resistive to desert climate (B). Gradually, strains are adapting to A climate in South America. The strains adapted to *“tropical-savannah” (Aw)* climate are a threat to all the tropical countries, which were initially less affected by COVID-19. Nevertheless, due to the uncertainty of COVID-19 data, the results should be carefully interpreted and should not be extrapolated to climate types and climatic conditions other than those analyzed here for the early evolution period. The study agrees that viruses are sensitive to their environment and respond towards naturally occurring abiotic factors such as temperature, latitude and humidity to sustain in different climate of the Earth, which also provides insights about seasonal variations possibly being a strong reason for the spread of other viral diseases as well. Here we showed a more refined description of genes based on phylogenetics and their distribution across different climates. This finer-grained analyses led to highly relevant insights on evolutionary dynamics of poorly understood SARS-CoV-2 genome and provides vital information about the direction of the spread and highlights vulnerable regions of Earth. Such inter-disciplinary studies will play an imperative role in designing antiviral strategies and taking pre-emptive precautionary measures to combat such a pandemic.

**Table 1:** SNPs representing virus cluster and their distribution across varied climates.

| Virus cluster | Nucleotide mutation | Amino acid mutation | Gene | Climate Zone | KCT | KC |
| --- | --- | --- | --- | --- | --- | --- |
| 1-22 | C8782T<br>T28144C | S2839<br>L84S | ORF1a<br>ORF8 | NTZ | Cfa | C |
| 5-6 | C29095T | F274 | N | NTZ | Cfa | C |
| 8-9 | T9477A<br>G25979T<br>C28657T<br>C28863T | F3071Y<br>G196V<br>D128<br>S197L | ORF1a<br>ORF3a<br>N<br>N | NTZ, TZ | Mix | C-A |
| 10-17 | C18060T | L1431 | ORF1b | NTZ | Cfa-Cfb | C |
| 12-17 | A17858G | Y1364C | ORF1b | NTZ | Cfa-Cfb | C |
| 13-17 | C17747T | P1327L | ORF1b | NTZ | Cfa | C |
| 20-22 | C24034T<br>T26729C<br>G28077C | N824<br>A69<br>V62L | S<br>M<br>ORF8 | NTZ | Cfa | C |
| 21-22 | T490A<br>C3177T<br>T18736C | D75<br>P971L<br>F1657L | ORF1a<br>ORF1a<br>ORF1b | NTZ,<br>NTSZ | Cfa | C |
| 23-25 | C6312A<br>C13730T<br>C23929T<br>C28311T | T2016K<br>L4489<br>Y789<br>P13L | ORF1a<br>ORF1a<br>S<br>N | NTZ, TZ,<br>SSTZ | Mix | Mix |
| 28-32 | G1397A<br>T28688C | D392G<br>L139 | ORF1a<br>N | NTZ | Mix | Mix |
| 33-43 | G26144T | G251V | ORF3a | NTZ | Cfa-Cfb | C |
| 37-39 | A2480G<br>C2558T | I739V<br>P765S | ORF1a<br>ORF1a | NTZ | Mix | Mix |
| 37-43 | C14805T | Y346 | ORF1b | NTZ | Cfa | C |
| 42-43 | T17247C | R1160 | ORF1b | NTZ | Cfb | B |
| 44-47 | G1440A<br>G2891A | G392D<br>A876T | ORF1a<br>ORF1a | NTZ | Cfb | C |
| 58-61 | C15324T | N519 | ORF1b | NTZ | Cfa-Dfb | C-D |
| 59-176 | C3037T<br>A23403G<br>C14408T | F924<br>D614G<br>P214L | ORF1a<br>S<br>ORF1b | NTZ | Cfa-Cfb-Dfb-Aw | C-D -<br>A |
| 59-125,<br>127-176 | C241T<br>C241T |  | Leader<br>seq. | NTZ | Cfa-Cfb-Dfa-Dfb | C-D-A<br>C-D |
| 66-68 | A26530G | D3G | M | NTZ | Cfc-Dfb | C-D |
| 70-71 | G4201T<br>C26527T | M1312I<br>A2V | ORF1a<br>M | NTZ | Cfa-Dwc | C-D |
| 80-115 | G28881A<br>G28882A<br>G28883C | R203K<br>R203K<br>G204R | N<br>N<br>N | NTZ | Cfa-Cfb-Dfb-Aw | C-D-A |
| 86-87 | C27046T | T175M | M | NTZ | Cfa-Dfb | C-D |
| 88-89 | C3373A | D1036E | ORF1a | NTZ | Dfb-Cfb | C-D |
| 105-107 | T29148C | I292T | N | TZ, STZ | Cfa-Aw | C-A |
| 106-107 | A27299C | I33T | ORF6 | NTZ, TZ | Cfa-Aw | C-A |
| 108-111 | C313T | L16 | ORF1a | NTZ, TZ | Cfa-Cfb-Aw | C |
| 113-115 | C4002T<br>G10097A<br>C13536T<br>C23731T | T1246I<br>G3278S<br>T4424I<br>T723 | ORF1a<br>ORF1a<br>ORF1a<br>S | STZ | Cfa-Cfb-Am | C-A |
| 116-125 | A20268G | L2167 | ORF1b | NTZ | Cfa-Cfb-Dfa-Dfb | C-D |
| 126-176 | G25563T | Q57H | ORF3a | NTZ | Cfa-Cfb-Dfa-Dfb | C-D |
| 126-130 | C18877T | L1704 | ORF1b | NTZ | Cfa-Dfa-Dcb | C |
| 131-135 | C2416T | Y717 | ORF1a | NTZ | Cfa-Dfa-Aw | D |
| 136-176 | C1059T | T265I | ORF1a | NTZ | Cfa-Cfb-Dfa-Dfb | C-D |
| 138-139 | C18998T<br>G29540A | A1744V | ORF1b | NTZ, TZ | Cfa-Am | C-A |
| 138-141 | C11916T | S3884L | ORF1a | NTZ | Cfa-Csb-Am | C |
| 143-147 | C27964T | S24L | ORF8 | NTZ | Cfa-Cfb-Dfa-Dfb | C-D |
| 148-149 | C11224T | V3653 | ORF1a | NTZ | Dfa-Dfb | D |
| 157-159 | G29553A |  |  | NTZ | Cfa | C |

**NOTE:** Virus clusters are named by Strain ID as depicted on the tree (Supplementary Table S1 and S2). Genomic coordinates in this study is based on reference genome<sup>20</sup>. The SNP positions are based on the reference genome. Nucleotide T represents nucleotide U in the SARS-CoV-2 RNA genome. Mutation at the protein level is not mentioned for the SNPs arising in the non-coding region. The amino acid position numbering is according to its position within the specified gene (CDS). In Climate zone column we have mentioned the major climate zone for the corresponding virus cluster<sup>18</sup>. KCT is Koppen's Climate Type and KC is Koppen's Climate columns display the main Koppen's climate in which most of the virus isolates of the corresponding virus cluster lie. 'Mix' implies no particular climate type is favored<sup>19</sup>.

## Potential caveats

We acknowledge that there are few caveats due to uncertainty in the COVID-19 data. The data from the tropical regions is limited because at the time of data collection (SARS-CoV-2 strains) from all over the world, the strains from the tropical countries were very limited, from few tropical regions strains were available (e.g., Ghana (Africa); India, Mexico, Nepal, Pakistan) but the data has been discarded due to the travel history of the strains, a large fraction of strains without travel history have large gaps in genomic sequences which were not suitable for the present study. Also, case history of each patient is not reported in the METADATA file as collecting all information from each patient is time-consuming. Hence, there are chances patients from whom these strains were isolated may have a migratory history. Data from different individual locations without travel history and large gaps in genomic sequences have been incorporated in this study. Due to these reasons, our analyses should be carefully interpreted and should not be extrapolated to climate types and climatic conditions other than those analyzed here for the early evolution period.

## Authors Contribution

Phylogenetic study and genomic analysis is carried out by P.B. GIS study and Koppen’s climate map interpretations is done by P.C.A. Statistical analysis is carried out by both the authors. Sequence alignment script was written by both the authors. Both authors have written, reviewed and edited the manuscript.

## Acknowledgement

We gratefully acknowledge the authors and originating and submitting laboratories of the sequences from GISAID’s EpiFlu (TM) Database on which this research is based. A table of the contributors is available in, Supplementary Table S1. We thank Prof. Raghavan Varadarajan, Prof. Raman Sukumar, Dr. Teena Jangid, and Chetankumar Jalihal of Indian Institute of Science for proofreading the article.

