## Supplementary Material for "Evolution and spread of SARS-CoV-2 likely to be affected by climate"

### Supplementary Tables and Figures

**Table S3:** List of description and criteria of different Koppen's climate types[19].

| S. No. | 1st | 2nd | 3rd | Description | Criteria |
|---|---|---|---|---|---|
| 1 | A | | | Tropical | $T_{cold} \geq 18$ |
| 2 | | f | | Rainforest | $P_{dry} \geq 60$ |
| 3 | | m | | Monsoon | Not (Af) and $P_{dry} \geq$ 100-MAP/25 |
| 4 | | w | | Savannah | Not (Af) and $P_{dry} <$ 100-MAP/25 |
| 5 | B | | | Arid | MAP < 10 ($P_{threshold}$) |
| 6 | | W | | Desert | MAP < 5 ($P_{threshold}$) |
| 7 | | S | | Steppe | MAP ≥ 5 ($P_{threshold}$) |
| 8 | | | H | Hot | MAT ≥ 18 |
| 9 | | | K | Cold | MAT < 18 |
| 10 | C | | | Temperate | $T_{hot}>10$ and $0 < T_{cold} < 18$ |
| 11 | | s | | Dry summer | $P_{sdry}<40$ and $P_{sdry} < P_{wwet}/3$ |
| 12 | | w | | Dry winter | $P_{wdry} < P_{swet}/10$ |
| 13 | | f | | Without dry season | Not (Cs) or (Cw) |
| 14 | | | A | Hot summer | $T_{hot} \geq 22$ |
| 15 | | | B | Warm summer | Not (a) and $T_{mon10} \geq 4$ |
| 16 | | | C | Cold summer | Not (a or b) and $1 \leq T_{mon10} < 4$ |
| 17 | D | | | Cold | $T_{hot}>10$ and $< T_{cold} \leq 0$ |
| 18 | | s | | Dry summer | $P_{sdry}<40$ and $P_{sdry} < P_{wwet}/3$ |
| 19 | | w | | Dry winter | $P_{wdry} < P_{swet}/10$ |
| 20 | | f | | Without dry season | Not (Ds) or (Dw) |
| 21 | | | A | Hot summer | $T_{hot} \geq 22$ |
| 22 | | | B | Warm summer | Not (a) and $T_{mon10} \geq 4$ |
| 23 | | | C | Cold summer | Not (a, b or d) |
| 24 | | | D | Very cold winter | Not (a or b) and $T_{cold} < -38$ |
| 25 | E | | | Polar | $T_{hot} < 10$ |
| 26 | | T | | Tundra | $T_{hot} > 0$ |
| 27 | | F | | Frost | $T_{hot} \leq 0$ |

**Acronym:** MAT, mean annual temperature; MAP, mean annual precipitation; $T_{hot}$, temperature of hottest month; $T_{cold}$, temperature of coldest month; $T_{10}$, number of months where temperature is above 10; $P_{dry}$, precipitation of driest month; $P_{sdry}$, precipitation of driest month in summer; $P_{wdry}$, precipitation of driest month in winter; $P_{swet}$, precipitation of wettest month in summer; $P_{wet}$, precipitation of wettest month in winter; $P_{threshold}$, logical (if 70% of map occurs in winter, then $P_{threshold}$ = 2(MAT), if 70% of map occurs in summer, then $P_{threshold}$ = 2(MAT)+28, else $P_{threshold}$ = 2(MAT)+14. Summer (winter) is defined as the warmer (cooler) six months period of ONDJFM (October-March) and AMJJAS (April-September).

**Table S4:** List of Koppen's Symbol for each climate type[19]:

| Koppen's symbol | Climate type |
|---|---|
| **Af** | Tropical rainforest climate |
| **Am** | Tropical monsoon climate |
| ***Aw or As*** | Tropical wet and dry or savanna climate |
| **BWh** | Hot desert climate |
| **BWk** | Cold desert climate |
| **BSh** | Hot semi-arid climate |
| **BSk** | Cold semi-arid climate |
| **Cfa** | Humid subtropical climate |
| **Cfb** | Temperate oceanic climate |
| **Cfc** | Subpolar oceanic climate |
| **Cwa** | Monsoon-influenced humid subtropical climate |
| **Cwb** | Subtropical highland climate or Monsoon-influenced temperate oceanic climate |
| **Cwc** | Cold subtropical highland climate or Monsoon-influenced subpolar oceanic climate |
| **Csa** | Hot-summer Mediterranean climate |
| **Csb** | Warm-summer Mediterranean climate |
| **Csc** | Cold-summer Mediterranean climate |
| **Dfa** | Hot-summer humid continental climate |
| **Dfb** | Warm-summer humid continental climate |
| **Dfc** | Subarctic climate |
| **Dfd** | Extremely cold subarctic climate |
| **Dwa** | Monsoon-influenced hot-summer humid continental climate |
| **Dwb** | Monsoon-influenced warm-summer humid continental climate |
| **Dwc** | Monsoon-influenced subarctic climate |
| **Dwd** | Monsoon-influenced extremely cold subarctic climate |
| **Dsa** | Mediterranean-influenced hot-summer humid continental climate |
| **Dsb** | Mediterranean-influenced warm-summer humid continental climate |
| **Dsc** | Mediterranean-influenced subarctic climate |
| **Dsd** | Mediterranean-influenced extremely cold subarctic climate |
| **ET** | Tundra climate |
| **EF** | Ice cap climate |

**Table S5:** Koppen climate type and their characteristics adapted by SARS-CoV-2[19].

| Koppen Climate Type | Characteristics |
|---|---|
| Humid-subtropical (Cfa) | The Koppen's symbol for the climate is 'Cfa', the 'C' of Cfa, denotes the temperature of the coldest month should be in between 0°C and 18°C, and the temperature of the hottest month is > 10°C. The 'f' is denoted for continuous rainfall throughout the year and 'a' means temperature of the hottest month temperature ≥ 22°C. |
| Marine-temperate (Cfb) | The 'C' and 'f' have the same definition as above, 'b' denotes hottest month's mean temperature < 22°C. |
| Mediterranean (Csa-Csb) | The 'C' has the same definition, 's' denotes dry winter, i.e., no rainfall during winters, 'a' and 'b' has similar meaning as above. |
| Humid-continental (Dfa and Dfb) | The 'D' represents, temperature of hottest month is greater than 10°C and the temperature of the coldest month is ≤ 0°C. 'f', has similar meaning and 'a', denotes temperature of hottest month is more than 22°C while 'b' denotes temperature of hottest month is < 22°C but temperature of 10 months is ≥ 4°C. |
| Tropical savannah (Aw) | The 'Aw' climate have monthly mean temperatures above 18°C in all months of the year with a typically a pronounced dry season, precipitation in the driest month is less than 60 mm. |

**Table S6: Contingency table for the chi-square test performed in this paper.**

| | Observed | Expected | *P*-value |
|---|---|---|---|
| **Environmental type** | | | |
| **Coastal** | 151 | 88 | 2.15E-21 |
| **Continental** | 25 | 88 | |
| **G1 variant group** | | | |
| **C** | 41 | 29 | 0.001625 |
| **Except C** | 17 | 29 | |
| **Russia** | | | |
| **G2** | 21 | 11.5 | 7.44E-05 |
| **G1** | 2 | 11.5 | |

#### Supplementary Figures

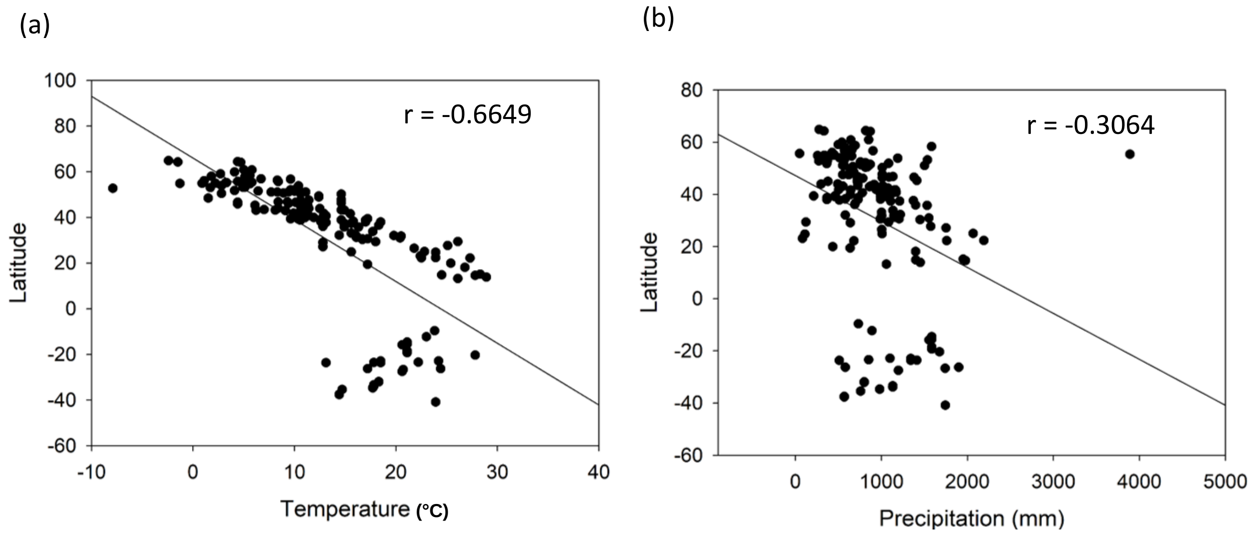

**Figure S1: Relationship between climatic variables.** Correlation between (a) latitude and temperature and (b) latitude and precipitation for SARS-CoV-2 strains (n=176) is estimated by calculating Pearson correlation coefficient (r).

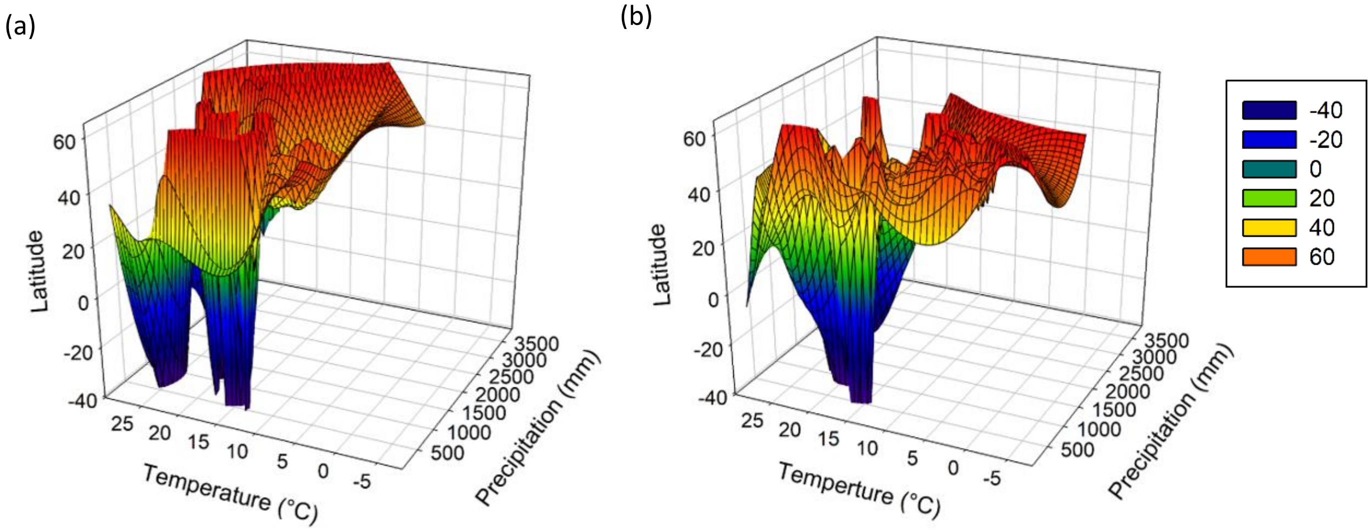

**Figure S2: Comparing climatic parameters such as latitude, temperature and precipitation for each SARS-CoV-2 isolate (n=176).** (a-b) Color code for the mesh plot is mentioned in the box and is according to latitude from which each SARS-CoV-2 strain was isolated. Latitude, temperature and precipitation values for each virus isolate are mentioned in Table S2. Relationship between latitude, temperature and precipitation for each SARS-CoV-2 strain belonging to (a) G1 variant group (Strain ID: 1-58) and (b) G2 variant group (Strain ID: 59-176).

(a)

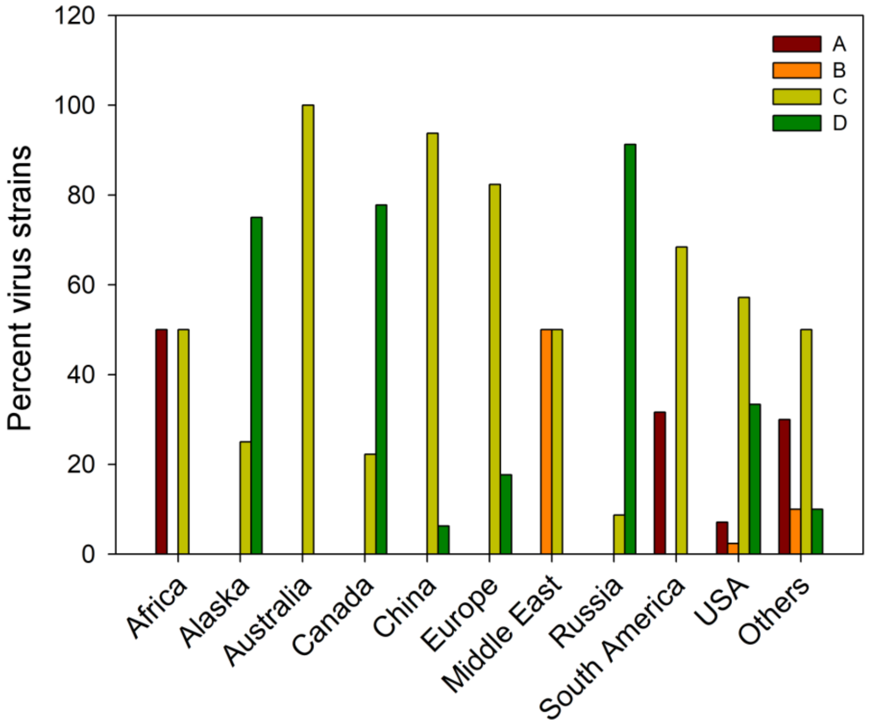

(b)

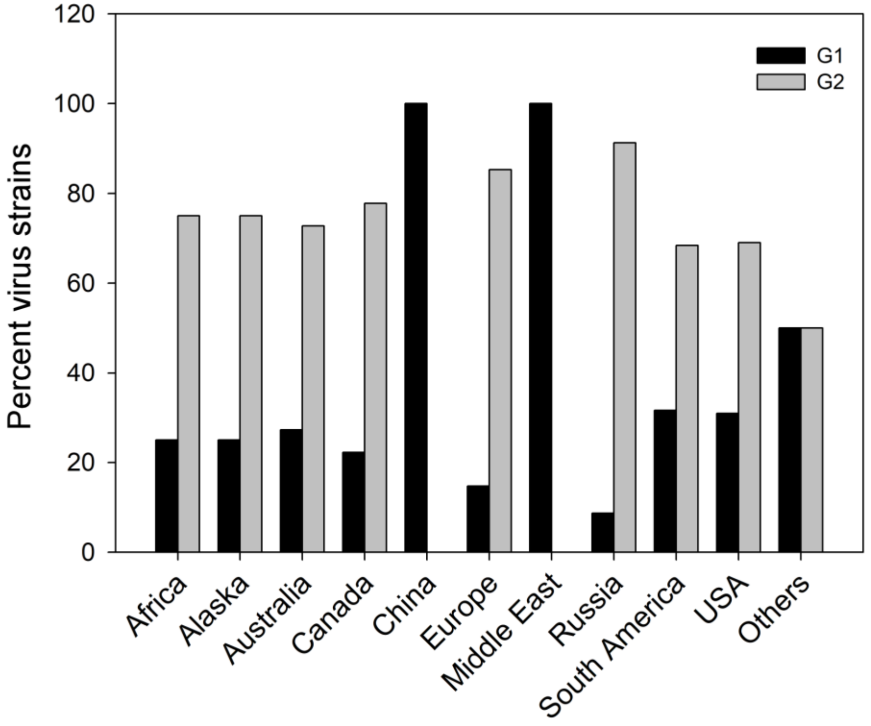

**Figure S3: Interpretation of Koppen's climate map country and/or continent wise:** Height of the bar represents percentage of virus strains (a) across A, B, C, and D Koppen's climate and (b) distribution of virus strains falling in G1 and G2 variant groups in major regions of the Earth. The G1 strains are abundant in China having specifically C climate. In Russia G2 strains are present in abundance having specifically D climate.

**Detailed interpretation and distribution of the strains on Koppen's map in major parts of the world (Figure 3, Supplementary Figure S3)**

**China:** The entire southeast China has "*humid-subtropical" (Cfa)* climate also referred in the text as Wuhan-type climate. This climate grades into the cold deserts in the north and the land is separated from ocean in the south which isolates SARS-CoV-2 strains (n=16) in '*Cfa'* climate. All strains (15/16) from China belong to G1 group (Figure 3). One G1 strain have appeared near to the C to D climate gradation near Beijing, suggesting G1 population can easily thrive in this climate (Chi-square test, $P < .001$).

**Europe:** The western coast of Europe consists of *"marine-temperate" (Cfb)* climate, a climate similar to Wuhan's climate i.e., '*Cfa*'. In UK, Spain, France, and Switzerland mainly *'Cfb'* climate persists, whereas Portugal has *"Hot-summer Mediterranean*" *(Csa)* climate. Towards west of Germany the '*Cfb'* climate dominates which grades into *"humid-continental" (Dfa-Dfb)* climate towards east which continues as a belt up to Japan. From the total strains (n=34), around 14.7% of G1 and 85.3% of G2 strains lie in Europe. All G1 strains (n=5) of Europe belong to C climate, of which 60% belongs to Cfb, 40% to Csa. Among G2 strains (n=29), 79.3% of G2 strains belongs to C climate (65.5% Cfa,10.3% Csa, and 3.4% Cfc) and 20.6% to Dfb climate type of D climate.

**Russia:** Majority (21/23) of strains (n=23) from Russia are present in the "*humid-continental" (Dfa-Dfb)* climate belt which begins from Germany and enters into the continent interior as a long (~8500 km) wide (~600-1700 km) belt, grading to *(Dwb-Dwc)* and tapering towards eastern side but continuing all along the southern boarders of Russia up to Japan. From Russia, ~8.7% of total strains (n=23), belongs to G1 and 91.3%

to G2. All G1 (2/23) strains are present in ‘*Dfb’* climate, one strain is present near the gradation of C to D (Strain ID: 4) climate and another (Strain ID: 31) in the interior of the continent. Of G2 strains, 91.3% of the strains are present in D climate (61.9% Dfb, 9.5% Dfa, 9.5% Dfc, 4.7% Dwb, 4.7% Dwc) and 9.5% in Cfa of C climate, suggesting a strong preference (Chi-square test, $P<.001$) of G2 strains towards D climate.

**North America:**

**USA:** Of all the continents North America USA has the most diverse climate, especially towards the western side[18]. The strains are mainly present in the eastern and western coasts of USA. The eastern coast of USA is one of the largest regions (~2.4 x$10^6$ $Km^2$) of the world having *“humid-subtropical” (Cfa)* climate (Wuhan’s climate). From the eastern coast, the strains enter into the continent interior through a long (~3800 km), wide belt (~600-1000 km) lying in the northern extremities of the USA, extending roughly in NW-SE direction initially following borders between USA and Canada and entering to Canada from southern side. This belt belongs to “*humid–continental” (Dfa-Dfb)* climate; is similar to that of Russia (mentioned above). The strains in the western coast of USA are aligned roughly parallel to the coastline and shows bulging towards the south (Figure 4), the western coast of USA has a bulged *“Hot-summer mediterranean*” *(Csa)* climate, which grades into *“humid-continental” (Dfa-Dfb)* towards its east. Between the western coast strains and eastern coast strains lies the cold desert, where SARS-CoV-2 strains are not present. In USA, ~31% of strains (n=42) belong to G1 while ~69% of the strains belongs to G2. Among G1 strains (n=13), 46.15% of strains belong to C climate (23% Cfa, 23% Cfb), 38.4% to D climate (15.3% Dfa, 23.07% Dfb) and 7.6% in both *“tropical-monsoon” (Am)* and “*cold-desert” (BSk)* climate. The G1 strains of D and A zones mainly lie near boundaries of C and D climate around the eastern and western coasts (Figure 4). Within G2 strains (n =29), ~62% strains lie in C climate (55% Cfa, 6.8% Csb), ~31% in D climate (20.6% Dfa, 10.3% Dfb) and remaining 6.8% are equally distributed in tropical climate (3.4% Am, 3.4% Aw).

**Canada:** The strains of Canada (n=9) are mainly distributed along the western coasts and towards the southern side. The western coasts of Canada have *"humid-subtropical" (Cfa)* climate and south of Canada has *"humid-continental" (Dfa-Dfb)* climate, which is an extension of '*Dfa-Dfb'* belt initiating from eastern side of USA near New York (mentioned above). 22.2% of the strains belong to G1 and 77.8% to G2. Within G1 (n=2), 50% strains belong to the '*Cfa'* and remaining to '*Dfb'*. Within G2 (n=7) variant group, ~14.2 % strains belong to *"marine-temperate" (Cfb)* and ~85.7% of strains belong to *"humid-continental" (Dfb)* climate. These small set of strains also show similar trend of G1 restricting to C climate and G2 prevalence in D climate.

**South America:** Majority (18/19) of South America's strains (n=19 of strains are present in the eastern coast of South America, The G1 population is concentrated in the Sao Paulo and one G1 strain (Strain ID: 3) is visible in the western coast of Chile, in both the places C climate is dominant, mainly *"humid-subtropical" (Cfa) and "marine-temperate" (Cfb).* Other than these two places the C climate is not present in the entire South America. The position and extent of South America in the globe is unique in itself as it connects the C climate with all A (tropical) climate through land. Both G1 and G2 strains are present in the C climate, however G2 strains have shifted towards the *"tropical-savannah" (Aw)* climate towards north, this shift is not visible towards the desert climate in south. Around ~31.5% of G1 strains and 68.4% of G2 strains are present in South

America. Within G1 (n=6), ~66% are present in the C climate (33.3% Cfa, 16.6% Cfb, 16.6% Csb), 33.3% in A climate (Aw). Within G2 (n=13), 69.23% of G2 strains are present in C climate (46.15% Cfa, 23% Cfb), while 30.73% of the strains are present in A climate (23% Aw, 7.6% Am).

**Africa:** Strains from Africa (n=4), are mainly (3/4) from G2 group. One G1 strains belongs to C climate (Cwb). Within G2, 66.66% of strains are present in A climate (Aw), and 33.33% in C climate (Csa).

**Australia:** All strains (n=11) from Australia are present either in the eastern or in the western coasts consisting of C climate. In the eastern coast the main climate is "*humid-subtropical" (Cfa)* and "*marine-temperate" (Cfb)* climate and western coast consists of *"Hot-summer mediterranean" (Csa)* climate of C climate. The rest of Australia has a desert climate. All strains from Australia are confined to this narrow belt of C climate. ~27% of the strains in Australia belong to G1 and ~73% of the strains belong to G2. Within G1 66.6% of strains are present in Cfa and 33.3% in Cfb. In G2, 50% of strains are present in Cfa, 25% in Cfb and 25 % in Csa climate type.

**Middle East:** The entire Middle East consists of B climate (desert). A very small portion of Middle East consists of C climate in the regions around Turkey. All strains (n=4) of Middle East belong to G1 group, among which 50% belongs to C climate (25% Csa, and 25% Csb) while the remaining are present in the "hot-desert" (BWh) climate.

**Japan:** Japan has mainly two climates i.e., *"marine-temperate" (Cfb)* towards south and *"humid-continental" (Dfb)* towards north. Both strain from Japan belongs to "humid-subtropical (Cfa)" climate.

#### South Asia and South Asian Islands

The G2 strains are present in India, Thailand and Vietnam, are mainly from "*tropical-savannah" (Aw)* climate, except strains from north-west India with a desert climate (BSh). The G1 strains are present in Philippines and South Korea are having "*tropical-savannah" (Aw)* and "*humid-continental"* (Dfa) climate respectively. The South Korea strain lies in the transition of *'Cfa'* climate (China) to '*Dfa'* climate (South Korea). Most of the strains in the South Asia and South Asian Islands belong to G2. Of total, around 80% of G2 and 20% of G1 strains are present in South Asia and South Asian Islands.
